# Double-membrane-spanning RNA export pores are a conserved feature in nidovirus replication organelles

**DOI:** 10.64898/2026.04.20.719621

**Authors:** Stanley Fronik, Georg Wolff, Ronald W.A.L. Limpens, Anja W. M. de Jong, Shawn Zheng, David A. Agard, Abraham J. Koster, Eric J. Snijder, Montserrat Bárcena

**Affiliations:** Section Electron Microscopy, Department of Cell and Chemical Biology, Leiden University Medical Center, Leiden, The Netherlands; Biohub, Redwood City, USA; Molecular Virology Laboratory, Leiden University Center for Infectious Diseases (LUCID), Leiden University Medical Center, Leiden, The Netherlands

**Keywords:** arteriviruses, coronaviruses, RNA replication, nonstructural proteins, double-membrane vesicles, cryo-electron tomography, subtomogram averaging

## Abstract

Upon infection, arteriviruses, coronaviruses, and other nidoviruses transform endoplasmic reticulum membranes into viral replication organelles. These include large numbers of double-membrane vesicles (DMVs) whose interior is considered the primary site of viral RNA synthesis. Early studies characterized nidovirus DMVs as sealed compartments, leaving it unclear how newly synthesized viral RNA could be exported to the cytosol. The discovery of DMV-spanning pore complexes in coronavirus-infected cells provided a plausible solution for this topological challenge. However, their structural organization, functional features, and evolutionary conservation across the nidovirus order, have remained unclear. Here, we investigated the macromolecular architecture of DMVs induced by two prototypic arteriviruses using cellular cryo-electron tomography. Despite the substantial evolutionary distance separating arteriviruses and coronaviruses, we observed DMV-spanning pore complexes with striking structural similarities to those previously described in coronaviruses. These pores appear to facilitate both export and encapsidation of viral RNA. In the absence of viral RNA synthesis, ectopic expression of the arterivirus transmembrane nonstructural proteins nsp2 and nsp3 sufficed to induce the formation of pore-containing DMVs. Together, our findings reveal the conservation of key structural features of DMV pores across two distantly related nidovirus families and support a central role for these pores in nidovirus replication.

## Introduction

The order *Nidovirales,* which currently comprises 14 virus families^1,2^([Taxon Details | ICTV](https://ictv.global/taxonomy/taxondetails?taxnode_id=202401821&taxon_name=Nidovirales)), was originally established to unify the distantly related *Arteriviridae* and *Coronaviridae*^3^, which share key similarities in the organization and expression of their polycistronic positive-sense RNA (+RNA) genomes^4^. While coronaviruses have gained worldwide attention in recent years as potentially lethal human pathogens – most notably following the 2020 pandemic caused by severe acute respiratory syndrome coronavirus 2 (SARS-CoV-2)^5^ - the arteriviruses identified to date infect only animal hosts.

The best-studied members of the arterivirus family^6,7^ include equine arteritis virus (EAV) and the porcine reproductive and respiratory syndrome viruses (PRRSV) 1 and 2^8^. Over the past decades, PRRSV in particular has had a profound global impact, causing substantial economic losses in the swine industry^9,10^.

Most families within the order *Nidovirales* possess relatively large genomes exceeding 25 kb, with the notable exception of arteriviruses, whose genomes range from ∼13 kb to 16 kb^6^. Despite this approximately two-fold size difference, the overall genome organization of arteriviruses is remarkably similar to that of other nidoviruses^4,11^ (Fig. 1). A defining feature is the presence of two large 5’-proximal open reading frames (ORFs 1a and 1b), which are translated into the replicase polyproteins (pp) 1a and 1ab, with the latter being a C-terminally extended version of pp1a that is produced via ribosomal frameshifting just upstream of the ORF1a stop codon. These two large polyproteins are proteolytically processed by internally encoded proteases to release the individual nonstructural proteins (nsps) (Fig. 1) that are the key orchestrators of the nidovirus replication cycle^12,13,6,14^.

**Figure 1.**
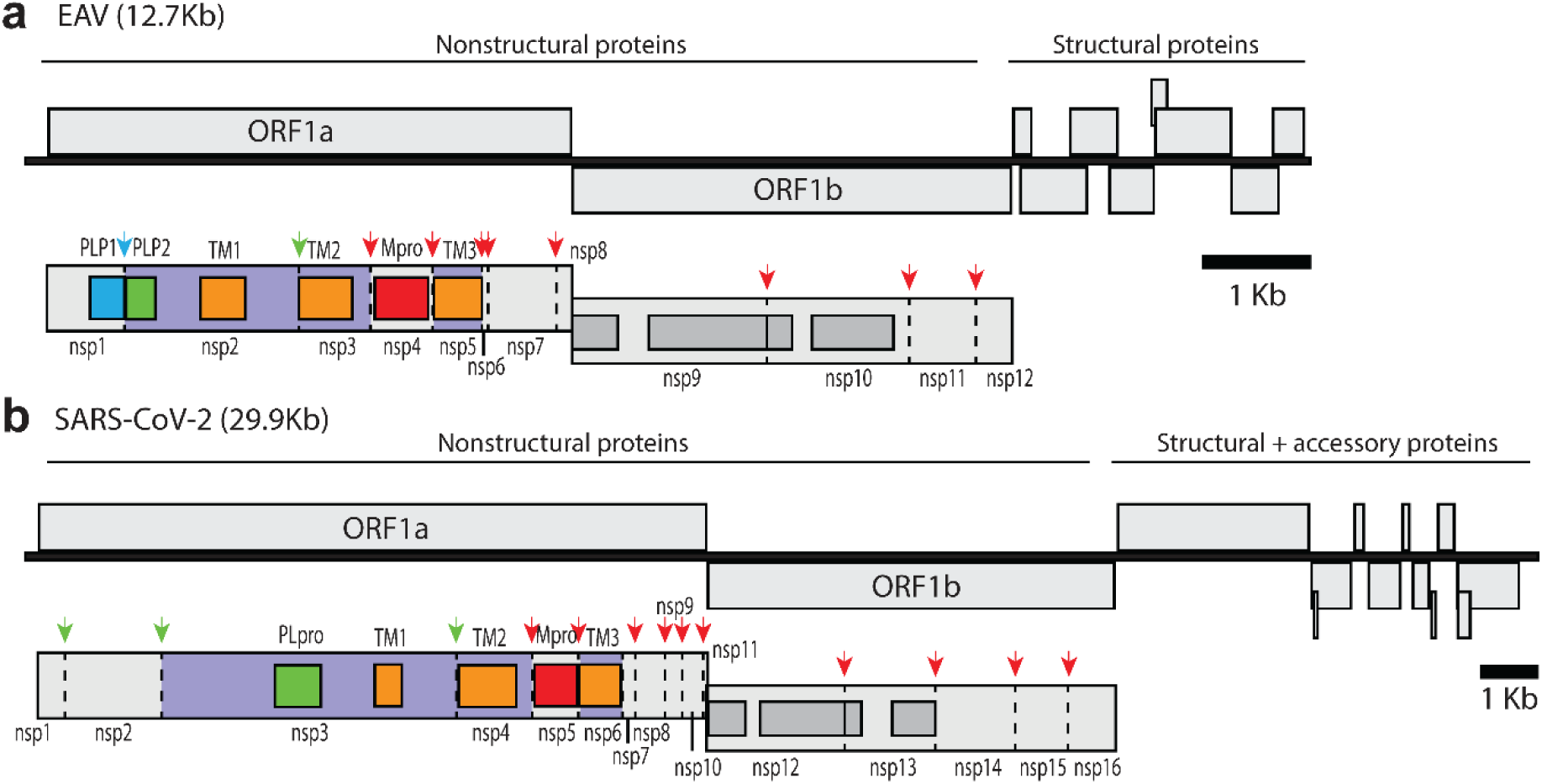
Comparison of the genome and replicase polyprotein organization of arteriviruses (EAV) and coronaviruses (SARS-CoV-2). Replicase gene and pp1ab map of EAV **(a)** and SARS-CoV-2 **(b)** are depicted using a 2:1 size ratio (EAV:SARS-CoV-2) for visualization purposes. A ribosomal frameshift signal just upstream of the ORF1a stop codon directs the extension of the ORF1a-encoded polyprotein (pp1a; not depicted) to produce pp1ab. Conserved ORF1b-encoded nidovirus domains implicated in viral RNA synthesis^11^ are depicted in grey, from left to right: NiRAN (Nidovirus RdRp-Associated Nucleotidyl transferase), RdRp, ZBD (Zinc-Binding Domain) and HEL (helicase). ORF1a-encoded proteases and their corresponding polyprotein cleavage sites are indicated by arrows in matching colors: EAV PLP1 (blue), PLP2 (green) and Mpro (red); SARS-CoV-2 PLpro (green) and Mpro (red). Transmembrane-domain-containing proteins (EAV nsp2, nsp3 and nsp5; SARS-CoV-2 nsp3, nsp4 and nsp6) are highlighted in violet, with the approximate location of their multi-spanning transmembrane domains (TM) shown in orange.

A hallmark of +RNA virus replication is the confinement of viral RNA synthesis to distinct, virus-induced membrane structures known as replication organelles (ROs)^15–19^. Nidoviruses remodel the host endoplasmic reticulum (ER) to generate an RO network of interconnected double-membrane structures^20–24^. The predominant components of this network are double-membrane vesicles (DMVs), which are widely regarded as the central hubs for viral replication. Replication transcription complexes (RTCs) – assemblies of nsps that contain the RNA-dependent RNA polymerase (RdRp) – associate with these compartments^25^, together with newly synthesized viral RNA^26^ and double-stranded RNA (dsRNA) replication intermediates^21,22,24,26^. Collectively, these findings support a model in which viral RNA synthesis occurs inside DMVs, providing a specialized microenvironment that facilitates replication while shielding RNA synthesis intermediates from innate immune detection^27^.

Nidovirus DMV formation has long been attributed to nsps that harbour three multi-spanning transmembrane domain regions, referred to as TM1, TM2 and TM3, which occupy the same relative position within the polyprotein of most nidoviruses^11,28^ (Fig. 1). These TM domains map to nsp3, nsp4 and nsp6 in coronaviruses, and nsp2, nsp3 and nsp5 in arteriviruses. Notably, the TM1- and TM2-containing proteins - coronavirus nsp3 and nsp4, and arterivirus nsp2 and nsp3 - are known to interact^29–32^. Although both these arterivirus nsps are considerably smaller than their coronavirus counterparts, their domain organizations share common features (Fig. 1). In both families, these proteins each harbour a luminal domain, flanked by transmembrane segments of the TM1 and TM2 domains, that likely mediates the interaction with their respective partner^29–31,33,34^ (Fig. 1). Taken together, these similarities point to conserved functional roles that extend to DMV formation, as this process can be triggered by co-expression of coronavirus nsp3 and nsp4, and arterivirus nsp2 and nsp3^29,31,32,35–37^. The role of the TM3-containing protein has so far been less clearly defined, but previous studies suggest it plays a more regulatory role in DMV formation and RO organization, in addition to modulating host cell metabolism^38–40^.

In conventional electron microscopy (EM) of resin-embedded samples, nidoviral DMVs appear as closed compartments without visible openings. However, the genomic RNA (gRNA) and subgenomic (sg) mRNAs synthesized inside DMVs must be exported to the cytosol for translation and, in the case of gRNA, packaging into progeny virions. Pioneering studies using *in situ* cryo-electron tomography (cryo-ET) revealed double-membrane spanning pores in the DMVs induced by the coronaviruses murine hepatitis virus (MHV) and SARS-CoV-2^41–43^. These pore complexes connect the DMV interior with the cytosol, thus providing a plausible gateway for viral RNA export.

Interestingly, the DMVs induced by ectopic expression of coronavirus nsp3 and nsp4 also contain DMV-spanning pores, indicating that these two proteins form the minimal machinery required to induce pore formation^32^. The structure of these pore complexes has recently been resolved at sub-nm resolution, using DMVs purified from cells expressing SARS-CoV-2 nsp3 and nsp4^31^. These complexes exhibit an overall six-fold symmetry and are composed of 12 copies each of nsp3 and nsp4, arranged into four stacked hexameric rings, with two distinct nsp3 hexamers forming the cytosolic crown and two distinct nsp4 hexamers forming the base of the pore^31^. A positively charged inner channel traverses the pore and contains three narrow constrictions ranging from 13 to 17 Å in diameter^31^, dimensions compatible with the passage of single- but not double-stranded RNA^44^. These architectural features are consistent with the proposed role of the DMV-pore complex in selective viral +RNA export.

While the nsp3-nsp4 expression systems have provided crucial insights into the architecture of DMV pores, they lack several key components required for the full RNA-synthesizing functionality of DMVs and their associated pore complexes, as observed in virus-infected cells. These absent elements include viral RNA, the TM3-containing nsp, and other viral proteins such as the RTC or nucleocapsid (N) protein, any of which may engage with the pore complex either transiently or stably. Notably, analysis of pore complexes formed in MHV-infected cells revealed additional densities at both the base and the cytosolic crown, the latter suggesting that the DMV-spanning pore may directly couple viral RNA synthesis and export with encapsidation^41^. Whether arteriviruses induce comparable DMV-spanning pore complexes has remained unknown. Previous ultrastructural studies of arterivirus-infected cells relied on resin-embedded, stained samples^22,35,45^, which offered detailed insights into DMV morphology but lacked the molecular resolution needed to resolve potential pore structures. Here, we employed cryo-ET to investigate arterivirus replication *in situ*, visualizing EAV- and PRRSV-infected cells under near-native conditions and in molecular detail. This analysis revealed that arteriviruses do indeed form DMV-spanning pore complexes and uncovered fundamental similarities with the structural organization of coronavirus DMV pores. Mirroring the coronavirus system, we identified nsp2 and nsp3 as the minimal components required for arterivirus DMV pore formation. Additionally, we observed numerous putative arterivirus encapsidation events in close proximity to DMV pores, supporting a model in which the pore complex serves as a central hub that couples selective viral RNA export with encapsidation. Altogether, our findings demonstrate that the conserved features of nidovirus ROs extend to molecular details such as the presence and architecture of DMV-spanning pores, suggesting that these structures may be preserved across the *Nidovirales* order and highlighting their likely essential role in the nidovirus replication cycle.

## Results

### Morphology of arterivirus ROs by cellular cryo-electron tomography

To investigate the morphology and organization of arterivirus ROs within their native cellular context, we employed cryo-focused ion beam milling^46^ to produce thin cryo-lamellae from vitrified Huh7 or MARC-145 cells infected with EAV or PRRSV, respectively. We selected relatively late infection timepoints, when ROs are highly abundant in the cytosol^22,45^. RO-rich regions from the cryo-lamellae were imaged and reconstructed by cryo-ET (Fig. 2).

**Figure 2.**
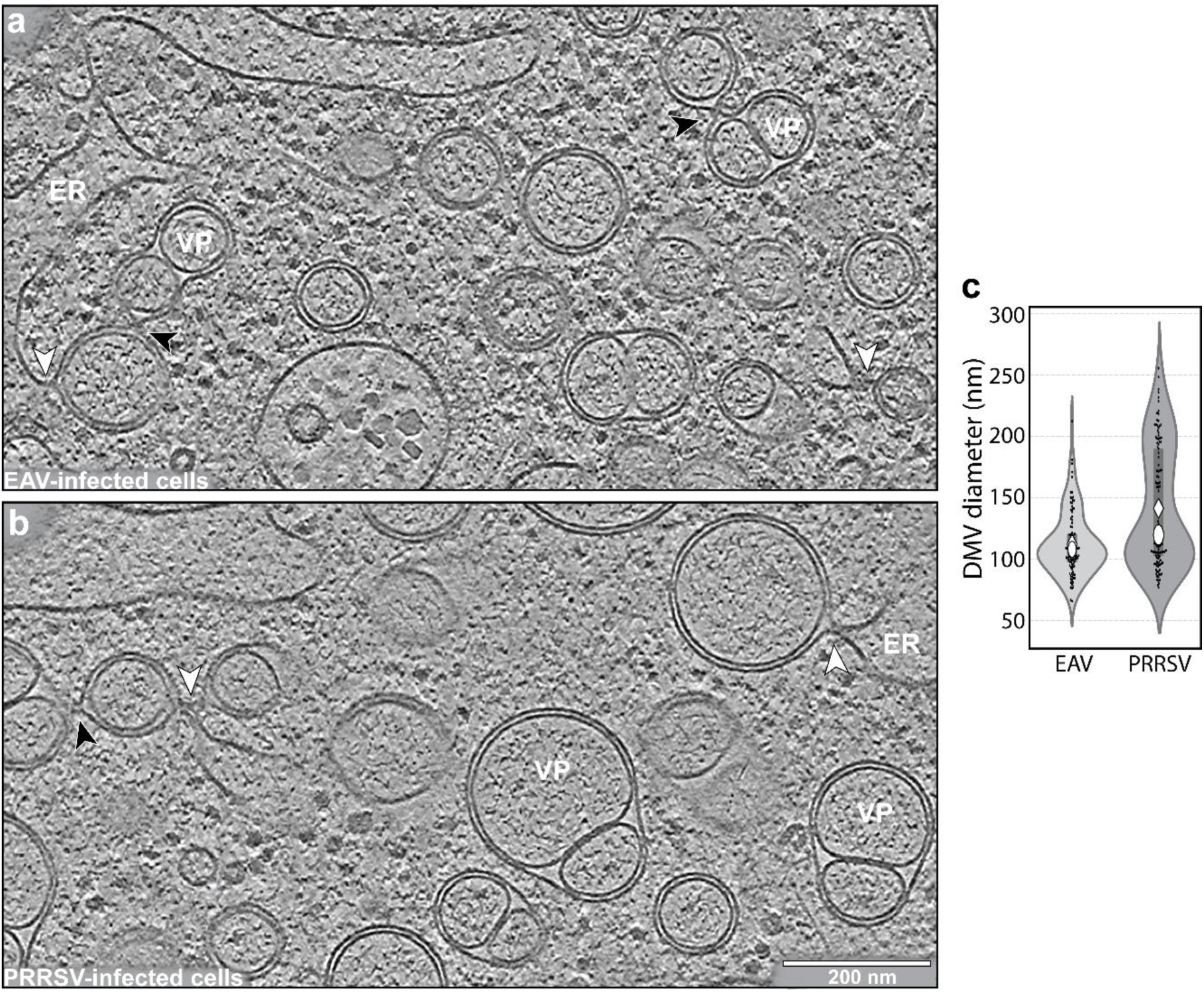
Cryo-electron tomography visualization of EAV and PRRSV replication organelles. Slices (13 nm thick) from **(a, b)** tomograms of cryo-lamellae from **(a)** an EAV-infected Huh-7 cell and **(b)** a PRRSV-infected MARC-145 cell. Selected vesicle packets (VP) and ER membranes (ER) are indicated. Neck-like connections between ER and DMVs (white arrowheads) as well as between adjacent DMVs (black arrowheads) were frequently observed. **(c)** Violin plots of arterivirus DMV diameters derived from the EAV and PRRSV data sets (EAV: *n* = 105 from 6 tomograms of 4 different cells; PRRSV: *n* = 121 from 9 tomograms of 3 different cells). White ovals denote medians and diamonds denote means; vertical lines within the violins indicate interquartile ranges. The two distributions differ significantly according to the Kolmogorov-Smirnov test (*p* < 0.001).

Arterivirus ROs consisted predominantly of DMVs as well as morphologically similar vesicles that appeared to arise from DMV outer membrane fusion. Although not previously documented in arterivirus-infected cells, they closely resembled the so-called vesicle packets (VPs) observed in coronavirus-infected cells at later stages of infection^21,42,43^. Arterivirus-induced VPs typically contained two – occasionally up to four – inner vesicles enclosed within a shared outer membrane (Fig. 2a, b). In line with previous reports^22,35^, arterivirus DMVs were substantially smaller than those observed in coronavirus-infected cells, with average diameters of 112 ± 26 nm for EAV and 141 ± 48 nm for PRRSV, compared to 200-350 nm for coronaviruses^21,23,41,42^. Size variability was particularly apparent for PRRSV, where large DMVs approaching coronavirus DMV dimensions were frequently observed, whereas EAV DMVs tended to be more uniformly small (Fig. 2c).

Imaging arterivirus DMVs in a frozen-hydrated state revealed distinct structural characteristics, closely paralleling the DMV features previously observed for coronaviruses by cryo-ET^41–43^. The inner and outer DMV membranes were separated by a uniform ∼4 nm gap, yielding an overall double-membrane thickness of ∼12 nm. This consistent and relatively narrow spacing suggests the involvement of proteins that bridge the space between the two membranes and maintain their close apposition. In line with such features, discrete mass densities were frequently observed within the intermembrane space (Supplementary fig. 1a, b). Distinct globular proteinaceous complexes could not be unambiguously identified within the interior of arterivirus DMVs, which – similar to their coronaviral counterparts - appeared filled with filamentous material. This filamentous content likely corresponds, at least in part, to dsRNA, a replication intermediate abundantly detected inside nidoviral DMVs by immunolabeling^21,22,24^.

ER membranes were observed in close proximity to nearly all DMVs in EAV- and PRRSV-infected cells. The DMVs and VPs exhibited neck-like connections between their outer membranes and the ER and/or adjacent DMVs (Fig. 2a, b; Supplementary Movie 1; Fig. 4a-d), together forming an extensive reticulovesicular network, a nidovirus RO hallmark^20–22,47^. Notably, free-floating DMVs - defined as DMVs clearly devoid of membranous connections to ER or other DMVs - were not detected in any of the datasets analyzed, suggesting that scission of DMVs from the reticulovesicular network is not a prevalent event in arterivirus-infected cells.

DMV biogenesis is thought to proceed through an enwrapping mechanism in which ER membranes first pair tightly to form zippered ER, which then curves and ultimately seals to generate the closed DMVs characteristic of nidovirus-infected cells^35,36,48^. Consistent with this model, we identified a few putative intermediates of DMV formation, including zippered ER with varying degrees of curvature and unsealed DMVs (Supplementary fig. 1 c, d, e). However, in line with previous reports, such intermediates were rare, supporting the notion that DMV formation in infected cells is a rapid process.

### Arterivirus-induced DMVs contain double-membrane–spanning pore complexes to which viral nucleocapsids associate

Since cryo-ET previously enabled the discovery of molecular pores in the DMVs induced by coronavirus infection^41^ or by ectopic expression of SARS-CoV-2 nsp3-nsp4^31,32^, we examined arterivirus-induced DMVs for comparable structures.In both EAV and PRRSV RO cryotomograms, pore complexes spanning the DMV double membrane were readily identified (Fig. 3a-d) and DMVs lacking detectable molecular pores were not observed in any dataset. The estimated number of pore complexes per DMV ranged from 1 to 8, with most DMVs containing 2 to 5 pores (Fig. 3b, d). Interestingly, our analysis suggests that the pore number does not correlate with DMV size (Fig. 3b, d), as also observed in the coronavirus nsp3-nsp4 expression system^32^.

**Figure 3.**
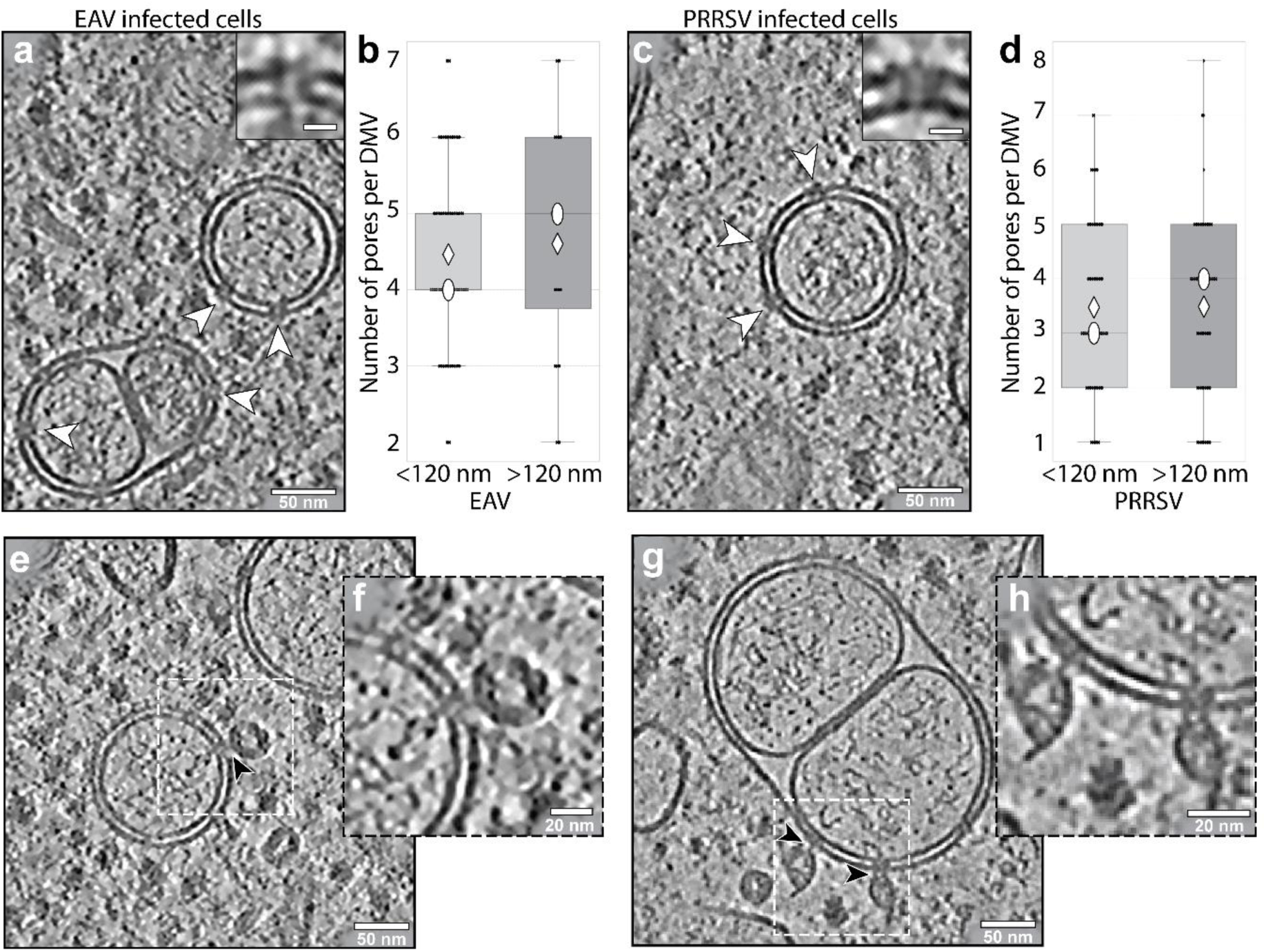
DMV-spanning pore complexes in arterivirus-infected cells and pore-associated nucleocapsids. Slices (13 nm thick) of **(a, c)** arterivirus-induced DMVs examples containing pore structures (white arrowheads) observed in **(a)** EAV-infected and **(c)** PRRSV-infected cells. Insets show a close-up view of individual pore complexes. Scale bar: 10 nm. **(b, d)** Boxplots showing the distribution of estimated pore numbers in arterivirus-induced DMVs measured in Fig. 2c. The average numbers of pores in DMVs classified as small (diameter < 120 nm, EAV *n* = 77, PRRSV n = 61) or large (> 120 nm, EAV *n* = 28, PRRSV *n* = 60) was similar. Consistent with this, the distributions for the two size groups were not significantly different **(b)** EAV *p* = 0.9625, **(d)** PRRSV *p* = 0.822, according to a Kolmogorov-Smirnov test. White ovals denote medians and diamonds represent means (see Materials & Methods for details on pore number estimation). **(e-h).** Nucleocapsids were frequently associated with pore complexes in DMV membranes (black arrowheads) in **(e, f)** EAV-infected and **(g, h)** PRRSV-infected cells.

The DMV-spanning pore complex is proposed to provide a gateway that enables newly synthesized viral RNA to exit the DMV interior and access the cytosol, where it is either translated into viral proteins or encapsidated by nucleocapsid (N) proteins^31,32,41,43^. Notably, RO-containing regions also featured additional macromolecular structures that appear to be involved in gRNA encapsidation and virus assembly. In both EAV- and PRRSV-infected cells, structures resembling viral nucleocapsids were frequently observed (Supplementary fig. 2a, b, c). The cytosol surrounding ROs contained numerous roughly spherical nucleocapsid assemblies with diameters of ∼37 and ∼ 42 nm in EAV- (Supplementary fig. 2a) and PRRSV-infected cells (Supplementary fig. 2b, inset), respectively. Their morphology and dimensions closely match those previously reported for purified PRRSV virions^49^, which consist of roughly spherical particles with an internal cavity and two concentric layers of N protein packaging the viral RNA between them, together measuring ∼10-11 nm in thickness with a subunit spacing of ∼7 nm^49^. In our tomograms, numerous particles appeared to be budding, or had already budded, into smooth-membrane compartments (Supplementary fig. 2a, b, c), likely derived from the ER or Golgi complex^6^, which strongly supports their identification as nucleocapsids. Association of nucleocapsids with DMV pores was consistently detected in both EAV- (Fig. 3e, f; Fig. 4a, e; Supplementary Movie 1) and PRRSV-infected cells (Fig. 3g, h). For example, nucleocapsid structures were associated with approximately 18% of DMV pores identified in PRRSV-infected cells (Fig. 3g, h), indicating a recurring spatial proximity between the two.

**Figure 4.**
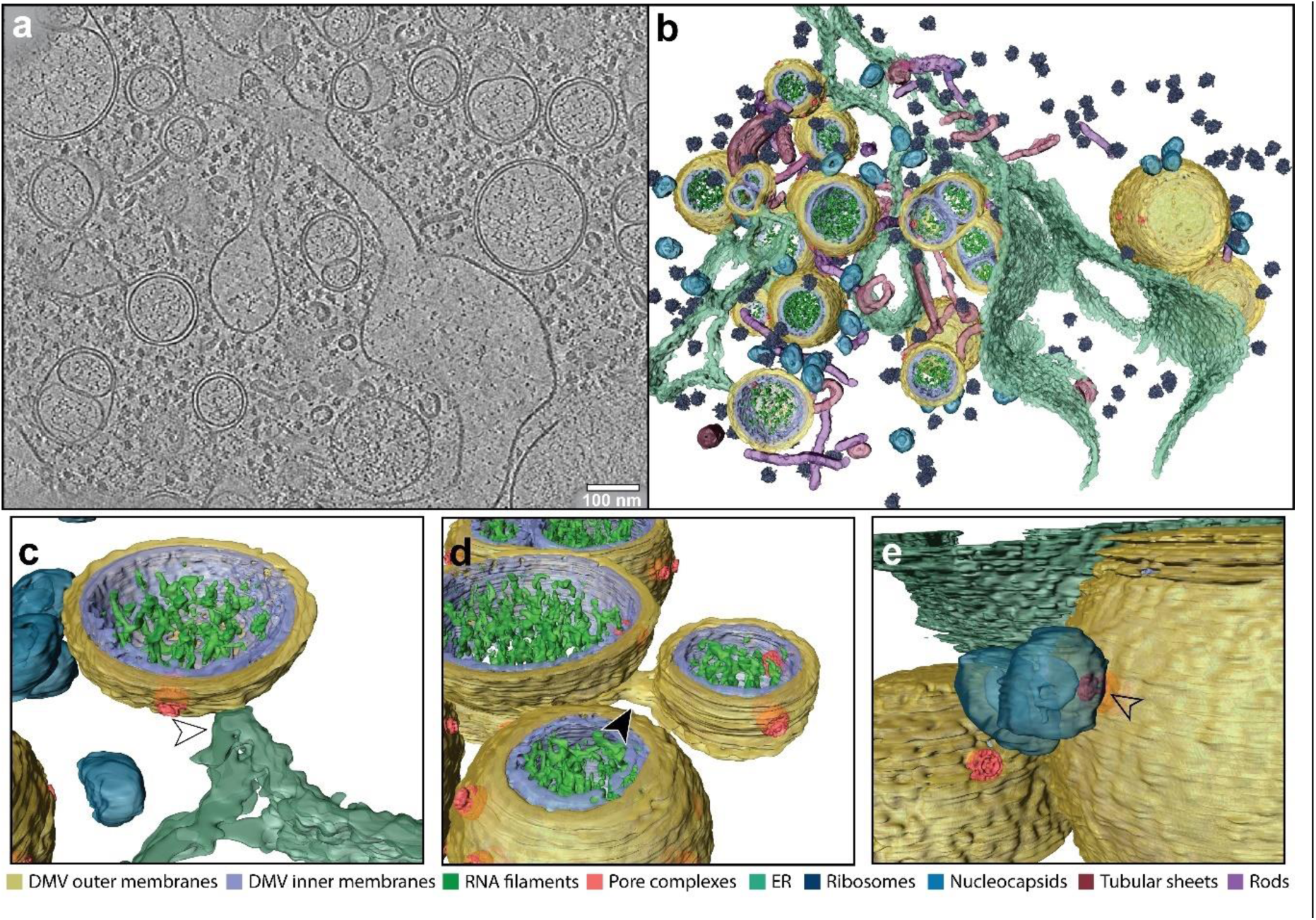
Segmentation model of EAV replication organelles. Slice (13 nm thick) from **(a)** a tomogram of a cryo-lamella from an EAV-infected Huh-7 cell. **(b)** 3D segmentation model of the tomogram displaying the following features: DMV outer membrane (dark yellow), DMV inner membrane (violet), putative RNA filaments (green), pore complexes (coral red), ER (teal green), ribosomes (indigo), nucleocapsids (blue), tubular sheets (Bordeaux red) and rods (purple). **(c, d, e)** Highlighted views from the segmentation model of **(c)** an ER-DMV membrane connection (white arrowhead), **(d)** a DMV-DMV membrane connection (black arrowhead), and **(e)** spherical nucleocapsid associated with the cytoplasmic face of the pore complex (for details about the pore complex architecture, see below). For visualization, EAV pore complexes and ribosomes were filtered to a resolution of 20 Å.

Despite the overall similarities, cytosolic EAV nucleocapsids were less abundant than in PRRSV-infected cells, and two additional distinct macromolecular assemblies were observed: tubular sheets and long, rod-like filaments (Supplementary fig. 2d, e; Fig. 4a, b; Supplementary Movie 1). The tubular assemblies have been reported previously in early 2D EM studies of cells infected with EAV or the related arterivirus lactate-dehydrogenase-elevating virus^50,51^. They were postulated to represent ribonucleocapsid (RNP) structures, as they labeled for N protein^22,51^ and were enriched in phosphate^22^, as expected for RNA-containing complexes. Regardless of their precise origin or function, it is striking that EAV nucleocapsids and these additional putative RNP-related assemblies were frequently associated with the cytosolic face of DMV pores, a pattern reminiscent of our previous observations in coronavirus-infected cells^41^. This spatial relationship supports the notion that the DMV pore complexes may act as central hubs for viral RNA encapsidation as newly synthesized gRNA exits the DMV interior.

### Architecture of the EAV DMV pore complex

Structural features of the coronavirus DMV pore complex that support its role in viral RNA export include a central channel that spans both membranes. It contains multiple constrictions that are too narrow to accommodate dsRNA but sufficiently wide to permit passage of ssRNA^31,44^. To investigate the architecture of arterivirus DMV-spanning pore complexes and assess whether their features parallel those of coronavirus DMV pores, we applied subtomogram averaging to our EAV pore dataset (Fig. 5). Without imposing symmetry, the resulting average revealed an intrinsic six-fold symmetry (Supplementary fig. 3c, d), consistent with the symmetry reported for coronavirus pore complexes^31,32,41^. Interestingly, additional densities were visible on both sides of the pore (Supplementary fig. 3e), as previously noted in subtomogram averages of coronavirus DMV pores reconstructed without imposed symmetry^41^.

**Figure 5.**
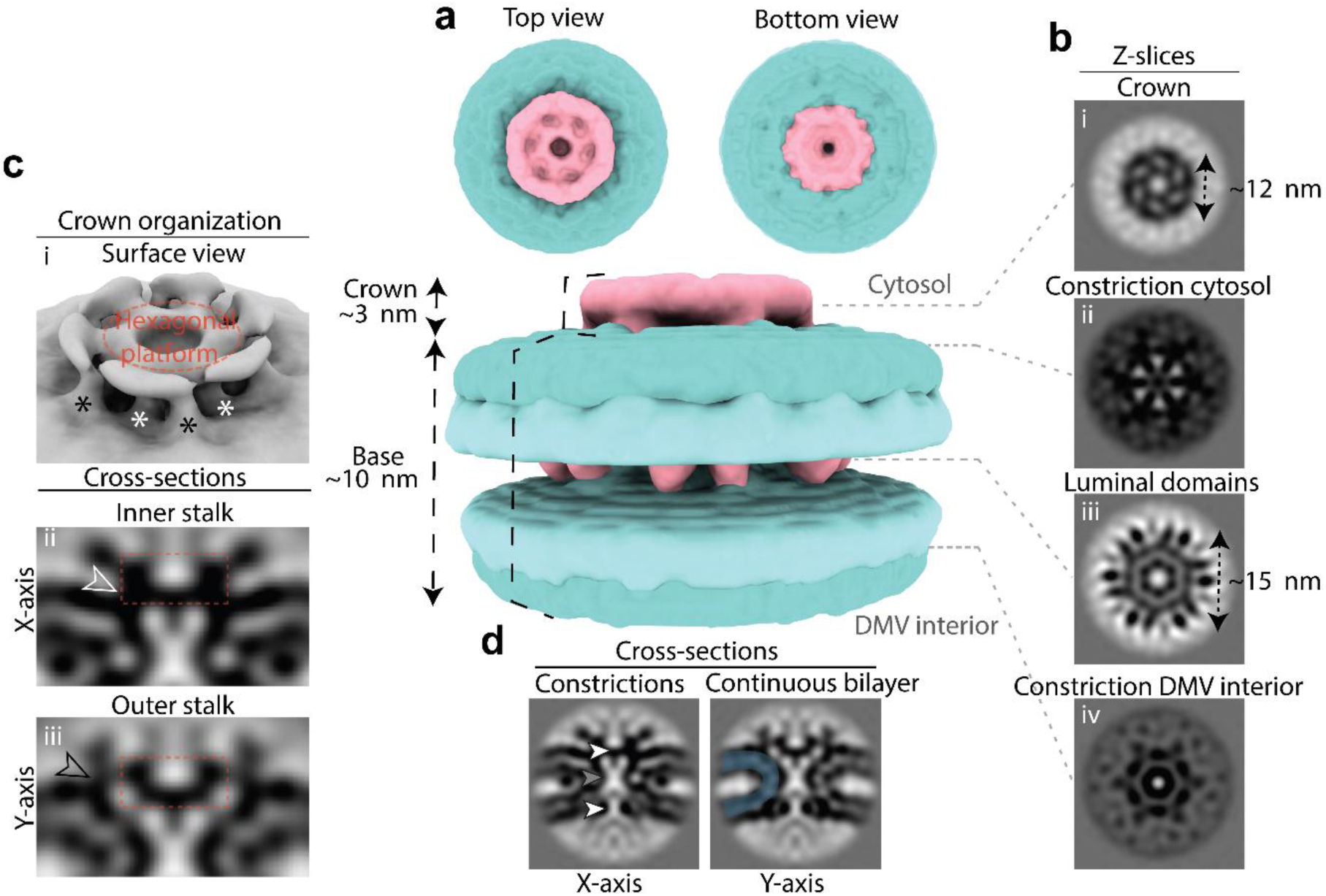
Subtomogram average of the EAV DMV pore complex. **(a)** 3D surface-rendered density map of the C6-symmetry EAV pore complex, showing top, bottom and side views. The membrane bilayer is shown in cyan and putative protein density in pink. **(b)** Cross sections through the EAV DMV pore complex at various positions along the z-axis: **(i)** At the crown-like structure facing the cytosol; **(ii)** At the constriction of the channel on the cytosolic side; **(iii)** At the luminal domains located in the intermembrane space; **(iv)** At the constriction facing the DMV interior. **(c)** Density map of the crown-like density with the hexagonal platform (red dotted circle or box), inner stalks (white asterisks) and outer stalks (black asterisks) highlighted. **(i)** High-threshold surface-rendering of the crown-like structure. **(ii)** x-axis cross section of the inner stalks (white outlined arrowhead). **(iii)** y-axis view of the outer stalks (black outlined arrowhead). **(d)** Cross-sections of the EAV pore complex, showing the channel. Putative constrictions are indicated by arrowheads, the cytosolic and DMV interior constrictions with white arrowheads and the central constriction by a grey arrowhead. The continuous membrane bilayer is traced in blue.

Subsequently, we imposed six-fold symmetry (C6), which yielded a reconstruction at a resolution of 20 Å from 3,450 particles (Supplementary fig. 3f). The final average revealed a pore complex approximately 13 nm in height, featuring a crown-like structure that extends ∼3 nm into the cytoplasm and having a maximum diameter of ∼15 nm within the DMV membrane (Fig. 5a-d). These dimensions are approximately twofold smaller than those of the coronavirus DMV pores resolved in MHV-infected cells and SARS-CoV-2 nsp3-nsp4 expressing cells, which measured ∼27 nm in height, including a ∼12 nm cytosolic crown, and had a membrane-embedded base with a diameter of ∼25 nm^31,41^. Consistent with this smaller size, we estimated a mass of ∼1.4 MDa for the arterivirus DMV pore complex at this resolution, which is significantly smaller than the estimated mass of the pore complexes from MHV-infected cells (∼3 MDa)^41^ and SARS-CoV-2 nsp3-nsp4 expression systems (∼2 MDa)^31^.

While the EAV DMV pore complex lacks the prominent cytosolic crown characteristic of its coronaviral counterpart, it nevertheless features a smaller crown-like structure that extends ∼3 nm into the cytosol and spans ∼12 nm in diameter (Fig. 5a, b(i)). When viewed from the cytosolic side, this crown consists of a hexagonal platform surrounded by a ring-like structure composed of six lobes (Fig. 5a, b(i), c(i)). Both elements are anchored to the outer DMV membrane by two distinct sets of six stalks arranged in an alternating pattern. An inner set of stalks originates from the hexagonal platform (Fig. 5c(i) white asterisks, (ii) white outlined arrow), while an outer set of stalks stems from the surrounding ring-like structure (Fig. 5c(i) black asterisks, (iii) black outlined arrow).

The widest part of the pore complex (∼15 nm in diameter) is located within the DMV intermembrane space (Fig. 5b(iii), h). At this level, twelve luminal domains assemble into a belt-like structure that encircles the pore complex and exhibits pseudo-12-fold symmetry (Fig. 5a, b(iii)). These luminal densities reside within the DMV intermembrane space and are flanked by the continuous, locally highly curved DMV membrane as it transitions from the outer to the inner membrane (Fig. 5d). This architecture is strikingly similar to that of the SARS-CoV-2 nsp3-nsp4 pore complex, in which the corresponding luminal densities have been shown to represent nsp3-nsp4 interaction sites^31^.

The complex is built around a channel that aligns with the six-fold symmetry axis (Fig. 5b, d). The channel diameter varies along its length and features three constrictions: one centrally located and two pronounced constrictions positioned at either end (Fig. 5b(ii),(iv), d). This architecture closely mirrors the three-constriction arrangement reported for the channel of the SARS-CoV-2 nsp3-nsp4 pore complex^31^. The cytosolic constriction marks the transition between the crown and the channel (Fig. 5b(ii)). At the present resolution, this region appears to be closed, suggesting that any opening at this site is smaller than ∼2 nm or may be occluded by translocating cargo. The constriction at the base of the channel, facing the DMV interior, measures approximately 2 nm in diameter (Fig. 5b(iv)), closely matching the narrowest constriction reported for coronavirus DMV pores in MHV-infected cells (∼2-3 nm) and in the SARS-CoV-2 nps3-nsp4 expression system (∼1.3 nm)^31,41^. A constriction of this size would permit the passage of ssRNA, but not dsRNA.

### Pore-containing arterivirus DMVs can form in the absence of viral RNA synthesis

To further investigate the requirements for DMV pore formation, we performed cryo-ET on lentivirus-transduced Huh-7 cells in which expression of an EAV nsp2-7 polyprotein could be induced^35^. This polyprotein included the three multi-spanning transmembrane nsps (Fig. 6a) as well as the nsp2 and nsp4 protease domains required for proper polyprotein cleavage. Inclusion of nsp6 and nsp7 ensured the generation of nsp5-7, which is a particularly abundant nsp5-containing cleavage product in EAV-infected cells^35,52^. Viral RNA synthesis is not possible in this expression system, as it lacks the viral gRNA as well as the RdRp subunit (nsp9) and other key RTC components. Nevertheless, nsp2-7-expressing cells were previously shown to generate DMVs closely resembling those formed during infection^35^. Our cellular cryo-ET analysis confirmed this close resemblance: smooth, spherical DMVs and vesicle packets were readily observed (Fig. 6b, c), along with characteristic neck-like connections to the ER and to adjacent DMVs (Fig. 6b, black arrowheads). As in infected cells, free-floating DMVs lacking membrane connections were not detected.

**Figure 6.**
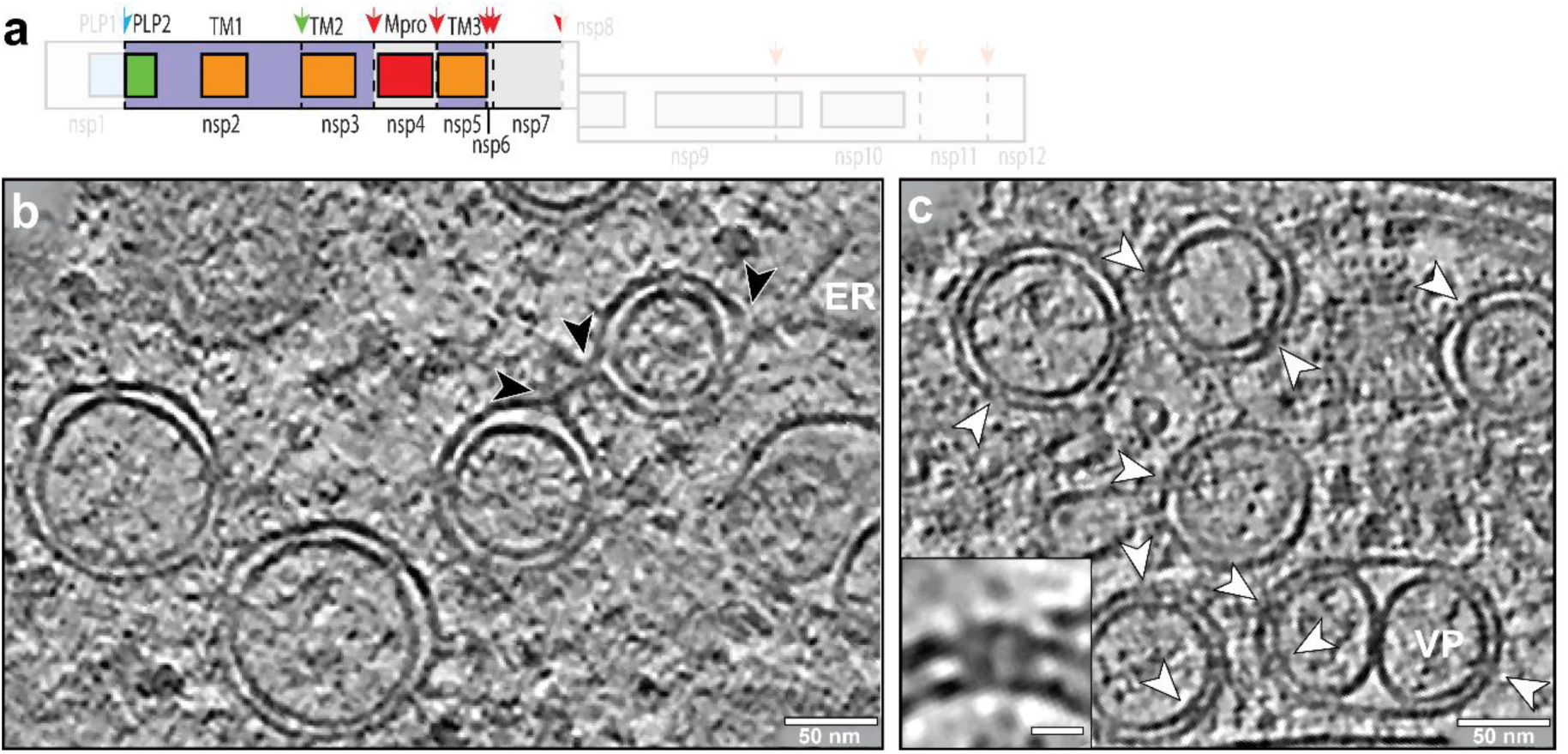
DMVs induced by EAV nsp2-7 expression contain pores complexes. **(a)** Schematic of the EAV nsp2-7 polyprotein, indicating transmembrane domains (orange) and protease cleavage sites for the PLP2 (green) and Mpro (red) proteases. Slices (13 nm thick) of **(b)** the overall morphology of nsp2-7-induced DMVs, which also formed VPs and established neck-like outer-membrane connections with adjacent DMVs and the ER (black arrowheads). **(c)** Visualization of pore structures (white arrowheads) embedded in the double membranes of nsp2-7-induced DMVs. A close-up side view (inset) highlights the resemblance with the pores present in DMVs in EAV-infected cells. Scale bar: 10 nm.

As expected, infection-dependent features were absent, such as fibrillar putative dsRNA within the DMV interior, nucleocapsids and related macromolecular assemblies (see above). Strikingly, our data revealed that the DMVs formed in this nsp2-7 expression system also contained pore structures closely resembling those observed in infected cells (Fig. 3a, 6c). Despite the absence of viral RNA synthesis, these pore complexes appeared comparably abundant (Fig. 6c). Within the limits of the current resolution, the pores displayed the same general organization as those in viral ROs, with an overall diameter of ∼15 nm and a central channel with a width of about ∼3 nm (Fig. 6c, inset).

### EAV nsp2 and nsp3 are sufficient to induce DMV and pore formation

Expression of a self-cleaving EAV nsp2-3 polyprotein was previously shown to induce arterivirus DMV formation^29,35^, analogous to the effects observed upon expression of the corresponding coronavirus nsp3-4 polyprotein. Given that coronavirus nsp3-4 expression sufficed to generate DMV pore complexes^31,32^, we next examined whether this minimal combination of multi-membrane-spanning nsps is likewise sufficient to generate arterivirus DMV pores. Notably, the DMVs formed in EAV nsp2-3 expressing cells exhibited pronounced morphological differences compared with DMVs formed upon nsp2-7 expression or during EAV infection (Fig. 3a, 6b, 7b). Their membrane curvature was less uniform, frequently resulting in oval rather than spherical DMVs (Fig. 7b). We frequently observed putative intermediates of DMV formation, such as zippered ER or double-membrane sheets that appeared to enwrap nascent DMVs but were not fully closed. Moreover, VPs were not detected in the nsp2-3 expression system, and both the average DMV diameter and overall size distribution were significantly larger than those observed upon EAV infection or nsp2-7 expression (Fig. 7d), consistent with previous findings^35^.

**Figure 7.**
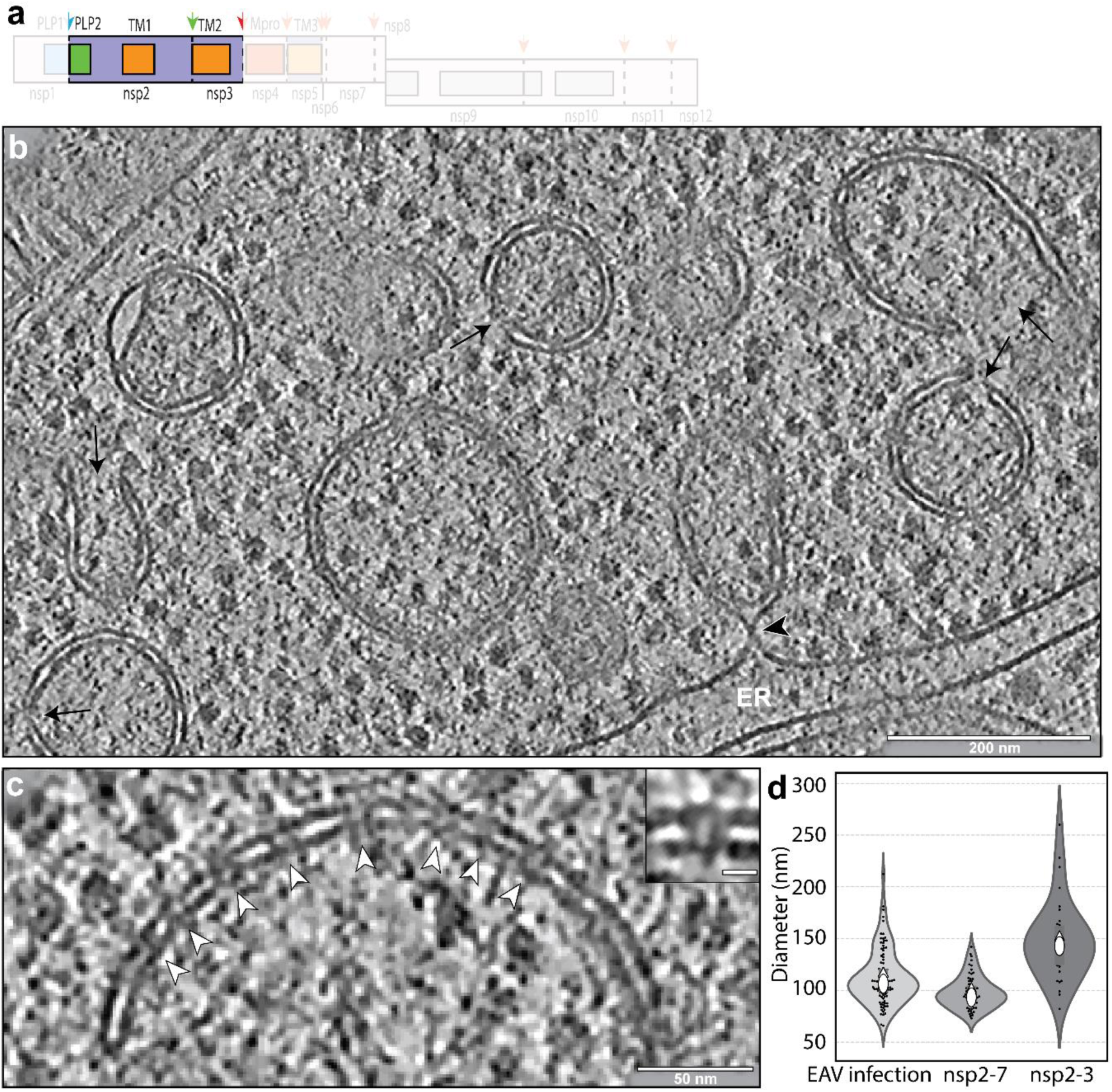
DMV pore complexes are formed in EAV nsp2-3-expressing cells. **(a)** Schematic of the EAV nsp2-3 polyprotein, indicating transmembrane domains (orange) and the PLP2 protease cleavage site (green arrow). Slices (13 nm thick) of **(b)** DMVs induced by nsp2-3 expression, showing irregular morphology, membrane openings (black arrows) and a neck-like connection to the ER (black arrowhead). **(c)** Pore complexes were frequently observed in close proximity to one another; this close-up reveals a row of pore complexes (white arrowheads) embedded in a double membrane. **(d)** Violin plots comparing DMV diameter distributions in EAV-infected Huh7 cells (*n* = 105 from 6 tomograms of 4 different cells) with those observed in Huh7 cells expressing nsp2-7 (*n* = 67 from 3 tomograms of 2 different cells) or nsp2-3 (*n* = 34 from 3 tomograms of 3 different cells) polyproteins. White ovals denote medians and diamonds denote means; vertical lines within the violins indicate interquartile ranges. The two distributions differ significantly according to the Kolmogorov-Smirnov test (*p* < 0.001).

Although the above differences suggest that DMV biogenesis was altered or impaired, pore structures were abundantly present in nsp2-3-induced DMVs. Morphologically, they also resembled those found in DMVs in EAV-infected cells (Fig. 3a, 6c, 7c, inset), featuring a central intermembrane platform that was ∼15 nm in diameter and a channel that was ∼3 nm wide. Pore complexes were particularly abundant in this nsp2-3 expression system and were especially frequent in larger DMVs, where they often occurred in close proximity to one another (Fig. 7c).

## Discussion

Replication organelles that support viral RNA replication are a hallmark of eukaryotic cells infected by +RNA viruses. Presumably, these virus-induced membrane structures provide an optimized microenvironment for viral RNA synthesis while shielding dsRNA replication intermediates from cytosolic innate immune sensors. Newly synthesized viral gRNA and sg mRNAs are subsequently exported to the cytosol for translation or packaging into progeny virions.

A common RO architecture among several groups of +RNA viruses consists of spherical membrane invaginations that maintain an open neck-like connection to the cytosol. Recent cryo-EM studies have identified elaborate ring-shaped protein complexes, or “crowns”, positioned at the cytosolic necks of these RO spherules (reviewed in ^19^). These viral protein assemblies are likely involved in coordinating RNA transport, and may also engage directly in RNA synthesis and mRNA capping^53–56^. For instance, the best-studied member of the nodavirus family, Flock House virus, induces RO spherules at the outer mitochondrial membrane that contain a distinct ring-shaped crown composed solely of protein A, organized as two stacked dodecamers^53,54,57^. Protein A is multifunctional, forming the crown structure while also harboring RNA synthesis and capping activities, suggesting that these functions are structurally integrated within a single macromolecular complex^57^.

A second major RO architecture comprises complex membrane networks that include multiple virus-induced membrane structures, among which DMVs are a recurrent and defining feature (reviewed in ^18,19,48^). Nidovirus-induced DMVs were initially considered sealed compartments, as no membrane discontinuities that could mediate direct export of newly synthesized viral RNA were observed. The discovery of a coronavirus DMV-spanning pore complex by cellular cryo-ET provided a compelling solution to this long-standing “transport riddle”^18,19,41^. Here, we investigated whether such pores represent a conserved feature across the nidovirus order by analyzing cells infected with arteriviruses, a family positioned at a considerable evolutionary distance from coronaviruses^11,58,59^. Despite the extensive sequence divergence and a more than twofold difference in replicase polyprotein size between arteri- and coronaviruses (Fig. 1), their ROs share multiple striking structural and functional parallels. These include the generation of large reticulovesicular networks of modified (double) membranes and the accumulation of numerous DMVs in infected cells. Our findings extend these similarities to the presence of DMV-spanning pore complexes, which likely provide an export pathway for viral RNA synthesized within the DMVs while also appearing to coordinate subsequent RNA encapsidation. Collectively, these data support the conclusion that the formation of pore-containing double-membrane compartments dedicated to viral RNA synthesis is a conserved feature of nidoviruses. This hypothesis can now be tested for representatives of additional families classified within this unique and evolutionarily diverse virus order.

Overall, the features of arterivirus ROs align well with previous studies describing an extensive reticulovesicular network of interconnected DMVs originating from ER membranes^21,22,35,41,42,45,60,61^. The mechanism underlying nidovirus DMV biogenesis remains to be fully elucidated, although studies using infected cells and heterologous nsp expression systems point to a model in which paired, zippered ER membranes progressively wrap to generate closed DMVs^18,31,35,36,48^. In agreement with earlier observations, the scarcity of putative intermediate structures in our samples further supports the notion that DMV formation is rapid. Examples of zippered ER and open DMVs - consistent with transient intermediates in the proposed enwrapping pathway - were only rarely observed in arterivirus-infected cells (Supplementary fig. 1). In contrast, such putative intermediates were highly abundant in cells expressing nsp2-3, indicating delayed, inefficient, or dysregulated DMV biogenesis in this system. Although nsp2-3 expression is clearly sufficient to induce DMV and pore formation - highlighting the central role of the nidoviral TM1 and TM2 domains in membrane remodeling^29,31,32,35,36,62,63^ - the resulting DMVs differed markedly from those formed during infection. Specifically, nsp2-3-induced DMVs were significantly larger than those in infected cells (Fig. 7d) and were frequently unsealed (Fig. 7b). This contrasts with results obtained using SARS-CoV-2 nsp3-4 expression, which yielded sealed DMVs that were significantly smaller than those formed during virus infection^31,32^, a difference attributed to the absence of viral RNA and other viral proteins^32^. Notably, this difference with the coronavirus data is partially mitigated in the EAV nsp2-7 expression system (Fig. 6b, 7d), where DMV diameters more closely match those observed in infected cells. In addition, nsp2-7-induced DMVs were predominantly spherical and morphologically similar to those formed during infection, while VP formation was also observed (Fig. 2a, 6b). These observations suggest that the TM3-containing subunit (nsp5 in arteriviruses) plays a more prominent role in DMV biogenesis than previously appreciated. TM3-containing nsps may directly modulate membrane curvature and/or act indirectly influence DMV formation by recruiting specific host proteins and/or lipids^38–40,64,65^.

Finally, in expression systems, the content of the DMV interior closely resembled the macromolecular landscape of the cytosol. In contrast, in both arterivirus- and coronavirus-infected cells, the DMV interior was dominated by putative (ds)RNA filaments (^42,48^ and this study). This difference suggests that DMV biogenesis in infected cells involves selective exclusion of cytosolic macromolecules, potentially ensuring preferential incorporation of viral RTC components and other factors required for viral RNA synthesis during DMV closure. A recent structural model offers a mechanistic framework for such selective recruitment^66^, proposing that pore complexes assemble from uncleaved replicase polyproteins, which would retain downstream RTC components at the RO membrane. As DMVs mature, gradual dimerization of Mpro may enable stepwise processing of the polyprotein into individual nsps, potentially occurring after DMV closure. Such a temporally regulated processing mechanism could facilitate coordinated RTC assembly within newly formed DMVs^66^.

As DMV pores appear to be a conserved feature of nidoviruses, it is reasonable to hypothesize that key aspects of their macromolecular architecture are similarly conserved. Our data demonstrate that nsp2 and nsp3, which contain TM1 and TM2 (Fig. 1), together constitute the minimal machinery required for arterivirus DMV pore complex formation. This parallels the role of nsp3 and nsp4 in coronaviruses^31,32^. However, both arterivirus proteins are less than half the size of their coronavirus counterparts, resulting in several pronounced structural differences between arterivirus and coronavirus DMV pore complexes. Specifically, the EAV pore is approximately twofold smaller in both dimensions (height and diameter) and estimated mass. Most strikingly, whereas coronavirus pore complexes possess a prominent cytosolic crown^31,32,41^, EAV and PRRSV pores display only a relatively small, crown-like density (Fig. 5a, c). This difference is readily explained by the marked size disparity between the crown-forming coronavirus nsp3 (∼2,000 amino acid residues in MHV and SARS-CoV-2) and the considerably smaller arterivirus nsp2 (572 residues in EAV and ∼1,100 residues in PRRSV).

Despite the absence of a prominent cytosolic crown, the EAV pore complex and its crown-like density appear to exhibit an organization strikingly analogous to that of the coronavirus DMV pore^31^. As observed in coronaviruses^31^, the EAV DMV pore is organized around a putative RNA export channel whose diameter varies along its length; it features three constrictions, including one at each end of the channel (Fig. 5b, d). The SARS-CoV-2 nsp3-4 pore complex comprises 12 copies each of nsp3 and nsp4, interacting via their luminal domains within the DMV intermembrane space, thus forming a belt-like structure with pseudo-12-fold symmetry^31^. A similar belt-like density is evident in the EAV pore complex (Fig. 5a, b(ii)), suggesting that it is composed of 12 copies each of nsp2 and nsp3. Our mass estimate of ∼1.4 MDa supports this interpretation, as it is compatible with the presence of 12 full-length copies of both nsp2 and nsp3. However, definite determination of the composition of the complex will require higher-resolution structural approaches, such as cryo-ET of purified DMVs^31^ or single-particle cryo-EM^67^ of solubilized, purified pore complexes.

In coronaviruses, both the N- and C-terminal domains of coronavirus nsp3 localize to the crown of the pore, where the PLpro domain on the external face interacts with the C-terminal CoV-Y domain, which extends from the internal face that generates the cytosolic constriction of the pore channel^31^. A comparable architectural arrangement may underlie the EAV pore complex. Here, six inner stalks emerge from the region that forms the cytosolic constriction of the channel and extend into a hexagonal platform that connects to six outer stalks via the ring-like structure (Fig. 5c). This remarkable structural parallel with the SARS-CoV-2 pore complex suggests that the region forming the cytosolic constriction and hexagonal platform corresponds to the C-terminal portion of EAV nsp2, whereas the lobes extending from the outer stalks and forming the ring-like structure likely represent the N-terminal PLP2 domain of nsp2.

Interestingly, data from the EAV nsp2-3 and nsp2-7 expression systems revealed that, as in coronaviruses, DMV pore formation does not require the presence of RTC components or viral RNA. In cellular cryo-ET studies of coronavirus ROs^41,42^, distinct densities that might correspond to RTCs were not conclusively identified in the DMV interior. In our symmetry-free subtomogram average of the EAV pore complex, however, we observed additional densities on both sides of the pore (Supplementary fig. 3e): one associated with the base of the pore complex within the DMV interior and another associated with the cytosolic crown-like density. These features were not resolved with sufficient clarity to allow definitive interpretation, suggesting either compositional heterogeneity or transient association. Notably, the density located at the base of the pore could potentially involve RTC components and/or viral RNA^41^, thereby providing a structural link between viral RNA synthesis within the DMV and RNA export through the pore channel. Definitive identification of additional components will require targeted follow-up experiments. It is also important to note that detecting the relatively small molecular assemblies predicted for nidovirus RTCs - for example, the proposed SARS-CoV-2 RTC composed of nsp7, 8, 9, 12 and 13, together measuring ∼300 kDa^68^ - may fall below the practical detection threshold of cellular cryo-ET^69^. Moreover, *in vitro* studies for coronaviruses have identified multiple polymerase-containing complexes with variable composition^68,70,71^, which may coexist and fulfil distinct functional roles^72^. Such compositional and conformational heterogeneity would further complicate their unambiguous identification within the crowded DMV interior. Accordingly, pinpointing the active RTC within the native nidovirus ROs remains a major unresolved challenge.

Although the presence of molecular pores in the DMVs strongly suggests a role in viral RNA transport across the double membrane, direct experimental evidence and mechanistic insight into the function of these pore complexes remain limited. In coronavirus-infected cells, nucleocapsid assemblies were frequently observed in close proximity to DMVs and, occasionally, in direct contact with DMV pores^41^. These observations led to a model proposing that DMV-spanning pores coordinate viral RNA export with encapsidation. Our current data from EAV- and PRRSV-infected cells provide strong additional support for this model. Nucleocapsid-like structures were frequently observed in direct association with DMV pores, consistent with immediate encapsidation of exported viral gRNA (Fig. 3e, f, g, h). Furthermore, the unidentified densities observed on the cytosolic side of the unsymmetrized EAV pore complex are compatible with this interpretation. Specifically, the density associated with the crown-like structure may correspond to N protein multimers, potentially in complex with viral gRNA. Although the precise functional roles and molecular identities of the additional putative RNP structures detected in EAV-infected cells - tubular sheets and rod-like assemblies - remain to be elucidated, their apparent interaction with DMV pores (Supplementary fig. 2d, e) further support this conceptual framework. A key unresolved question in arteriviruses and other nidoviruses concerns how gRNA is selectively encapsidated over sg mRNAs. Only a minority of the +RNAs synthesized by nidoviral RTCs are full-length gRNAs, and only a subset of these are ultimately destined for packaging into progeny virions. In contrast, the vast majority of viral +RNAs consist of sg mRNAs to be translated into viral structural or accessory proteins.

The selective encapsidation of gRNA may be coordinated at the level of the DMV pore complex, potentially in conjunction with progressively increasing concentrations of N protein in the cytosol of the infected cell. However, the molecular basis for such selectivity remains unclear. Arterivirus nsp2 has been reported to interact with N protein^73^, while coronavirus nsp3 harbors an N-terminal ubiquitin-like 1 (UBL1) domain that likewise interacts with N protein^74–76^. In addition, also the 5′ untranslated region of SARS-CoV-2 RNA was recently shown to interact with nsp3^77^. These findings may represent the first indications for a mechanistic link between RNA export through the pore and subsequent encapsidation by N protein. Nevertheless, they do not fully explain how gRNA is discriminated from sg mRNAs, which thus remains an intriguing open question. In summary, our findings establish DMV pore complexes as a conserved feature of nidovirus replication. These structures likely serve as central hubs that couple viral RNA synthesis and export to encapsidation. Given this pivotal role, DMV-associated pore complexes represent attractive targets for antiviral intervention.

## Acknowledgements

We thank Frank Faas, Roman Koning and other members of the Electron Microscopy section (LUMC), members of the Lamers group, Cell and Chemical Biology (LUMC), Sander Roet, Wouter Beugelink and other members of the Structural Biochemistry group (Utrecht University) for advice on image processing and helpful discussions. We are grateful to Ying Fang (University of Illinois, Urbana-Champagn, USA) for providing the PRRSV isolate SD01-08 and for helpful discussions. We thank Diede Oudshoorn and Marjolein Kikkert (LUMC) for sharing the EAV nsp-expressing cells and Jessika Zevenhoven-Dobbe, Linda Boomaars-van der Zanden, Emmely Treffers, Nina de Beijer and other colleagues of the LUMC Molecular Virology group for their assistance and helpful discussions. The cryo-ET data was collected at the Netherlands Centre for Electron Nanoscopy (NeCEN) with assistance from Wen Yang, Christoph Diebolder and Ludovic Renault. Access to NeCEN was made possible through financial support from the Dutch Roadmap Grant NEMI (NWO grant 184.034.014). This work was supported by the Netherlands Organization for Scientific Research (grant NWO-OCENW.M.21.339).

## Author contributions

The conceptualization of the study was carried out by S.F., G.W., A.J.K., E.J.S. and M.B. Investigation was undertaken by S.F., G.W., R.W.A.L.L., A.W.M.J. and M.B., while formal analysis was conducted by S.F., G.W. and M.B. Visualization was carried out by S.F., G.W., R.W.A.L.L. and A.W.M.J. Resources were provided by S.Z., D.A.A., A.J.K. and E.J.S. Data curation was performed by S.F., G.W., A.J.K. and M.B. The original draft of the manuscript was prepared by S.F., G.W., A.J.K., E.J.S. and M.B., and all authors contributed to reviewing and editing the manuscript. Supervision and funding acquisition was provided by A.J.K., E.J.S. and M.B.

## Materials and Methods

### EAV and PRRSV infection

EAV infections were performed in Huh-7 cells, which were maintained as described previously^35^. Sixteen hours prior to infection, cells were seeded at a density of 1 × 10^4^ cells per cm^2^ into culture dishes containing EM grids and incubated at 37 °C. Cells were infected with a cell-culture-adapted derivative of the EAV Bucyrus isolate^78^ at a multiplicity of infection (MOI) of 50 and incubated at 39.5 °C. At 9 h p.i., cells were plunge frozen for analysis.

PRRSV infections were performed in MARC-145 cells, which were maintained at 37 °C and 5% CO_2_ in Dulbecco’s modified Eagle’s medium (DMEM) supplemented with 8% fetal bovine serum (FBS), 100 U/mL penicillin, and 100 μg/mL streptomycin. Cells were seeded at a density of 4 × 10^4^ cells per cm^2^ and reached full confluence after three days, the state in which they are most susceptible to PRRSV infection. Cells were infected with PRRSV genotype 1 isolate SD01-08 [GenBank accession #DQ489311^79^] at an MOI of 2 and incubated at 37 °C. Since confluent cell layers form a cohesive mass that precludes efficient vitrification of the entire cellular volume by plunge freezing, infected cells were trypsinized at 17 h post infection (p.i.), diluted 1:10 in cell culture medium, and reseeded into 10-cm^2^ culture dishes containing EM grids. At 24 h p.i. cells were plunge frozen for analysis.

### Cells stably expressing EAV nsps

Lentivirus-transduced Huh-7 cells expressing the EAV nsp2-3 or nsp2-7 polyproteins^35^ were cultured in selection medium containing 500 µg/ml G418 (PAA Laboratories) and 12.5 µg/ml blasticidin-S (PAA Laboratories). Seven hours prior to induction, cells were seeded at a density of 8 × 10^3^ cells per cm^2^ into culture dishes containing EM grids. Expression was induced for 24 h using 1 µg/ml tetracycline (Invitrogen), after which cells were plunge frozen.

### EM sample preparation for cellular cryo-electron tomography

R1/4 gold Quantifoil grids (200 mesh, Quantifoil Micro Tools) were cleaned overnight on a chloroform-soaked filter paper in a sealed glass dish, dried, glow-discharged, and placed in 10 cm^2^ culture dishes containing PBS. Grids were UV-sterilized for 30 min in a laminar-flow cabinet. After PBS removal, cells were seeded onto the grids at the indicated densities. For vitrification, grids were plunge-frozen in liquid ethane using a Leica EM GP automated plunge-freezer (Leica Microsystems). Prior to freezing, grids were blotted from the reverse side for 10 s, and subjected to a 1 s post-blot delay, 95% humidity and 37 °C (EAV) or 39 °C (PRRSV). Grids were stored in liquid nitrogen until further processing. For cryo-lamella preparation, grids were transferred to an Aquilos cryo-focused ion beam scanning electron microscope (FIB-SEM; Thermo Fisher Scientific), and cryo-lamellae generated as described previously^80^.

### Cellular cryo-electron tomography

Tilt-series were acquired on a Titan Krios (Thermo Fisher Scientific) electron microscope operated at 300 kV and equipped with a Gatan BioQuantum energy filter (Gatan) and either a Gatan K3 summit direct detection device (the latter was used for part of the EAV dataset). Movie frames were collected in counting mode with zero-loss imaging and a 20-eV slit. Pixel sizes were 3.28 Å. Defocus ranged from −4 to −8 µm. Dose-symmetric tilt series^81^ were collected with SerialEM software^82^, and bidirectional tilt series with FEI Tomo4 (K2). The total electron dose per series was 120-140 e^−^/Å^2^, distributed over a tilt range of 100°- 120° with 2 or 3° increments. The number of high-quality tilt series collected used for analysis was 29 for EAV-infected cells, 20 for EAV nsp2-3 expressing cells, 6 for EAV nsp2-7 expressing cells and 14 for PRRSV-infected cells.

### Tomographic data processing

Motion correction, tilt-series alignment, reconstruction, and local CTF correction were performed in Aretomo3 v2.2.2^83^. Both motion correction and tilt series alignment were executed with 7×5 patches (local alignment). The tomograms were reconstructed using WBP (weighted back projection) with local CTF correction. For visualization of specific tomographic slices, tomograms were denoised with nonlinear anisotropic diffusion^84^ in IMOD^85^.

### Subtomogram averaging

Pore complexes were selected from tomograms of EAV-infected cells reconstructed from K3 camera data. final tomograms were reconstructed at bin4 using WBP with local CTF correction and imported into Scipion3 v3.8.1^86^, and denoised using cryoCARE v0.3.0^87^. Particles were manually picked in denoised tomograms (3,985 particles for EAV), using Eman2 tomo v2.99.55^88^. Particle coordinates were converted into Relion 4-compatible particles.star format^89^, using the prepare data functionality for Relion 4 particle lists in Scipion3. To prepare the tilt series alignment information generated by AreTomo3 for subsequent subtomogram averaging with Relion 5, a (slightly) modified version aretomo3torelion5 (https://github.com/Phaips/aretomo3torelion5/) was used. To convert the Relion 4 compatible particle coordinate list particles.star to Relion 5 format, a (slightly) modified version (https://github.com/SBC-Utrecht/SBCscripts/blob/main/rel4_particles_to_rel5/convert_rel4_rel5.py) was used. Prior to subtomogram averaging with Relion 5, the values of first Euler angle (in-plane rotation of the particle, rlnAngleRot in Relion) in the particle list were randomized. All subtomogram averaging steps were performed in Relion 5^90^. The initial model for the EAV pore complex was generated de novo using a 300 Å spherical mask, 100 iterations, a regularization parameter (T) of 4, and no imposed symmetry (C1). To center the resulting model, C20 symmetry was applied. The first 3D auto refine job was executed with global search settings. All subsequent 3D auto refine jobs in the workflow were performed with local search settings. After the first 3D auto refine job, the assigned orientation of the particles was inspected using ChimeraX^91^ and ArtiaX^92^. This revealed that some particles had been incorrectly assigned as top or bottom views, even though only side views of pore complexes had been manually picked. These particles, with a tilt angle value <60° or >120°, were removed from subsequent alignment steps. Refinements proceeded from binned to unbinned data with CTF refinement and Bayesian polishing. For EAV, the average was generated by 3D refine jobs with local search, sequentially with binned 4x, 3x 2x and 1x pseudo-subtomograms, followed by a single refinement iteration (CTF refine and Bayesian polishing) and a final 3D refine job at 1x binning, which resulted in a 26 Å resolution map (C1). To generate a C6-EAV pore complex, C6 symmetry was imposed after the first refinement iteration, followed by a 3D refine job and a second refinement iteration (CTF refine and Bayesian polishing), and a final 3D refine job at 1x binning, which resulted in a 20 Å resolution map (C6). For visualization, EAV C1 and EAV C6 pore maps were subjected to a post processing job, sharpened with a B-factor of −200 Å^2^, and filtered to their final resolution. Subtomogram averaging of ribosomes from EAV--infected cells was performed to confirm the correct handedness of the tomograms. Four tomograms were selected for template matching with PyTom_tm v0.12.0^93^, using an *in-situ* subtomogram averaging derived ribosome density map (EMD-15879)^94^ as template. A total of 2,000 particles were refined in Relion 5^90^. yielding a final average at 15 Å resolution.

### Segmentation model

Segmentation was performed in Amira (Thermo Fisher Scientific). The 3D surface renderings of the DMVs and other organelles were segmented following a semiautomatic segmentation protocol previously described^95^.DMV pore positions and orientations were placed manually using ChimeraX^91^ and ArtiaX^92^. The header voxel spacing of the pore averages (EAV), was increased 1.5 times in the Z dimension only, using IMOD^85^ for visual purposes. The ribosome average obtained from EAV-infected cells as described before was filtered to 20 Å and used to indicate the position and orientations of some ribosomes present in the region. Ribosome and pore complex averages were positioned on the particle coordinates with their corresponding Euler angles using ChimeraX^91^ and ArtiaX^92^ and imported into Amira for placement in the segmentation.

### Data analysis and statistics

DMV pore positions and DMV centers were annotated in IMOD^85^. For a DMV center, the central point at the DMV equator was selected. The pore positions were annotated in the center of the inter-membrane platform. The annotated coordinates were extracted and used for the quantifications of DMV sizes and number of pores per DMV. DMV radius was calculated as the average pore-to-center distance plus 6 nm (distance from pore center to the cytosolic face of the outer membrane). For EAV nsp2-3 samples, only spherical DMVs were included. To estimate the total number of pores per DMV for DMVs not fully contained in the lamella we assumed: (I) that DMVs are spherical and (II) that the pore density is uniform across the DMV surface. Accordingly, the fraction of DMV surface present in the-lamella was calculated, and the number of detected pores was then extrapolated to the full DMV surface. Even though VPs contained abundant molecular pores, for simplicity, we excluded vesicle packets. Statistical comparisons used Kolmogorov-Smirnov tests to assess statistically significant differences between distributions. The whisker plots (Fig. 3b and d) show in boxes the 2^nd^ (bottom) and 3^rd^ (top) quartiles, the whiskers extending to the minimum and maximum values measured, while outliers that exceeded Q2 or Q3 by a distance of 1.5 times the interquartile range are depicted as dots. Outliers were included in the calculation of the medians (middle lines) and means (crosses). For the violin plots, white ovals denote medians and diamonds denote means. vertical lines within the violins indicate the interquartile ranges.

### Data and code availability

The subtomogram average map of the EAV DMV pore complex (C6 symmetry) has been deposited in the Electron Microscopy Data Bank (EMDB) under accession code EMD-57473.

