## Supplemental Figures 1-2-3 for "Double-membrane-spanning RNA export pores are a conserved feature in nidovirus replication organelles"

### Supplementary material

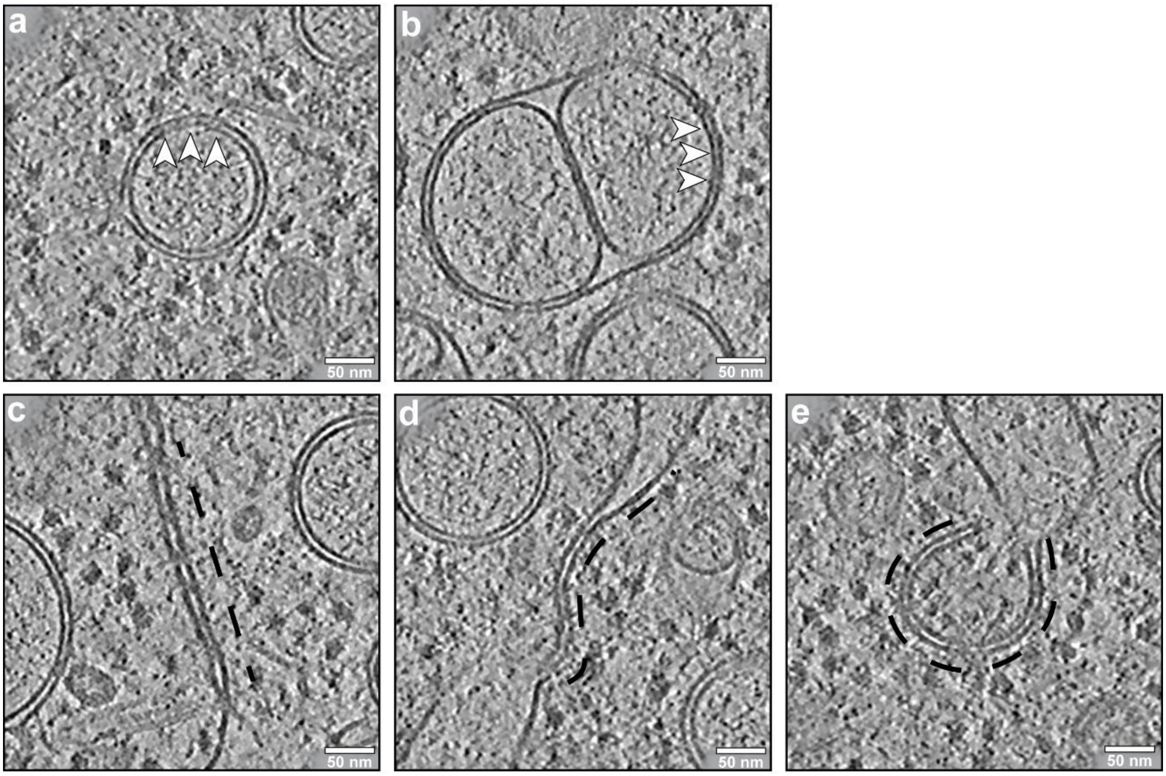


**Supplementary figure 1 – DMV bridging densities, ER membrane zippering and possible intermediates in DMV biogenesis.** Slices (13 nm thick) of **(a, b)** bridging densities spanning the membranes of **(a)** EAV- and **(b)** PRRSV-induced DMVs (white arrowheads). **(c-e)** Putative intermediates in DMV biogenesis **(c, PRRSV)** straight zippering of ER membranes **(d, EAV)**, curving of the zippered ER **(e, EAV)**, and an unsealed DMV. The membrane curvature of zippered ER is outlined with a black dotted line.

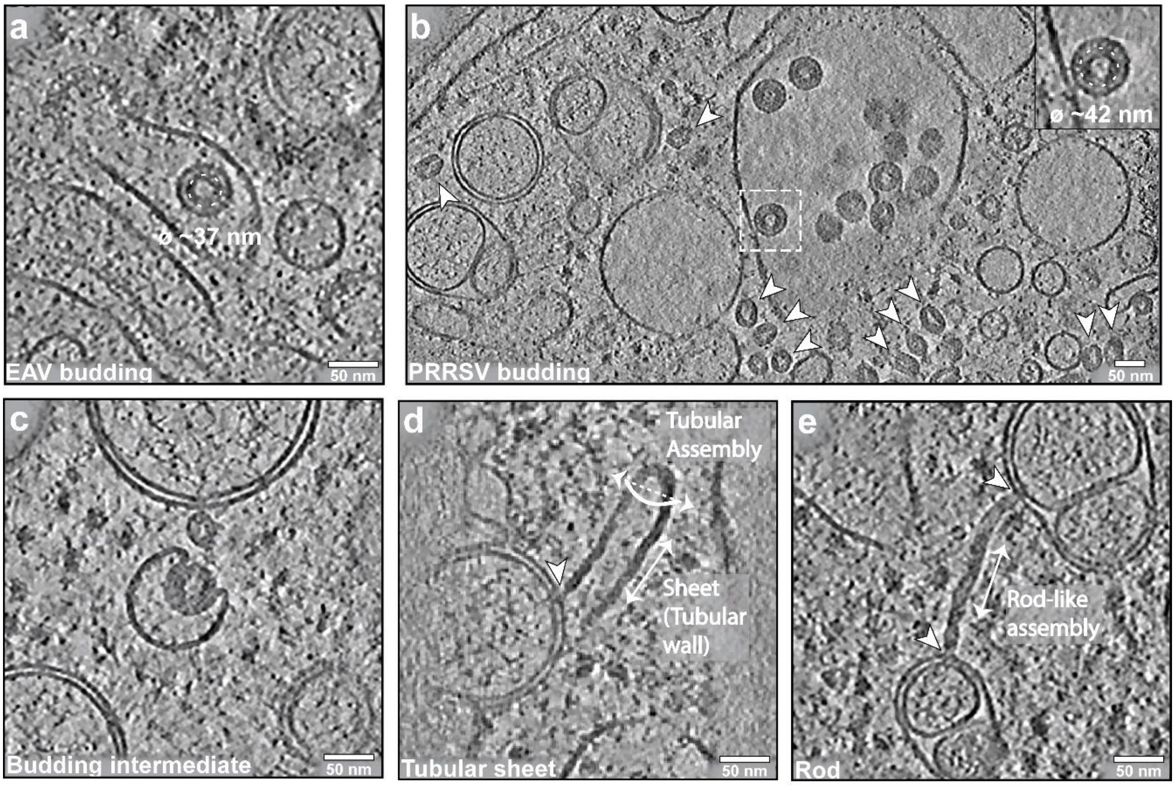


**Supplementary figure 2 – Arterivirus budding events and putative tubular RNP sheets and RNP rods observed in EAV-infected cells.**

Slices (13 nm thick) of **(a-c)** regions containing DMVs in **(a)** EAV- and **(b, c)** PRRSV-infected cells, with roughly spherical nucleocapsid structures (~37 and ~42 nm in diameter for EAV and PRRSV, respectively) frequently observed in budding virions. **(d)** Sheets that form larger, tubular assemblies were regularly found in EAV-infected cells, occasionally associated with DMV pores (black arrowhead). **(e)** Rod-like molecular structures were abundant in RO areas as well, some exceeding lengths of more than 150 nm, often associated with DMV pores (black arrowheads), extending into the cytosol perpendicular to the DMV membrane.

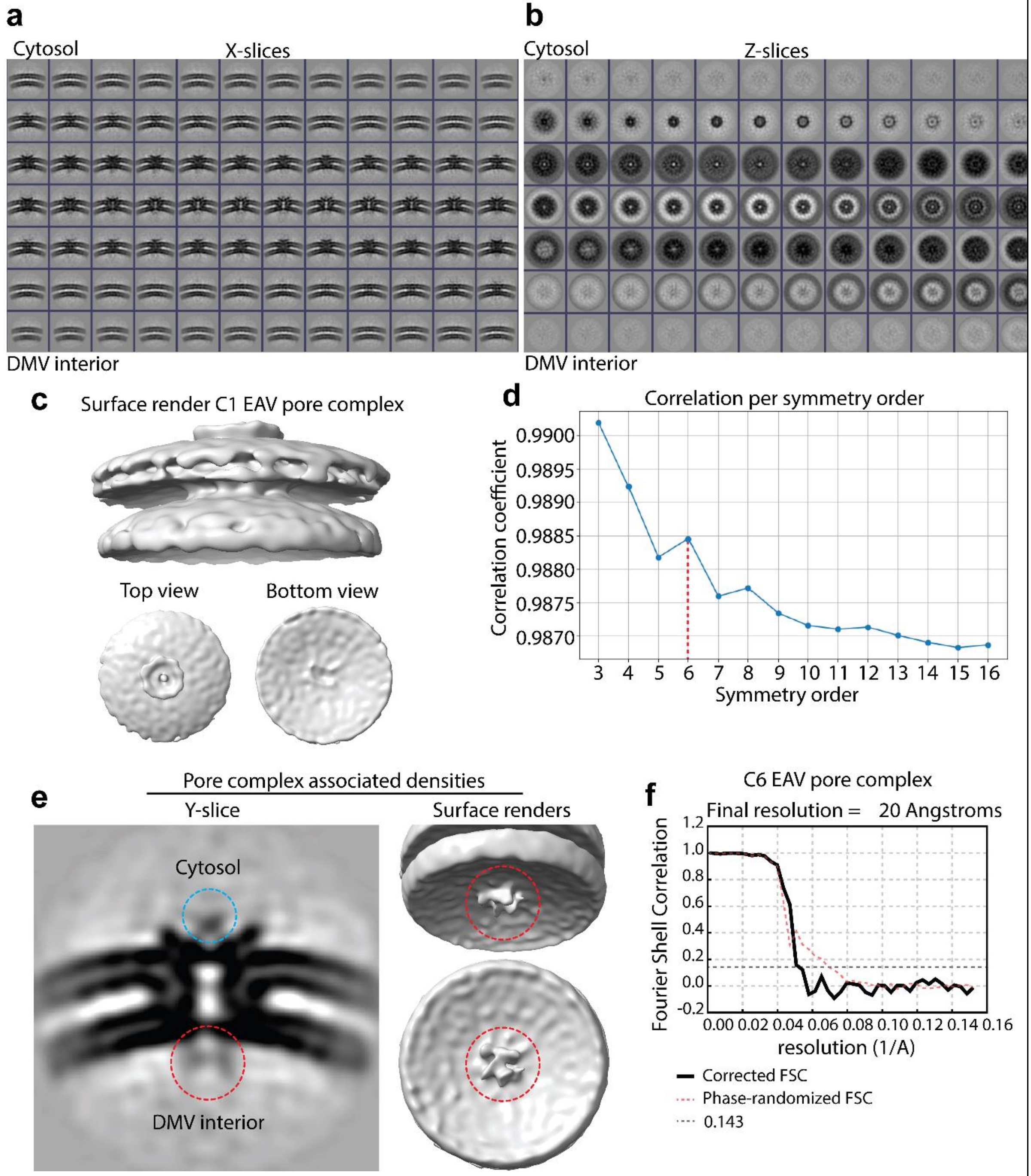


**Supplementary figure 3 – Subtomogram averaging details of the EAV DMV pore complex.**

**(a, b)** Gallery of the C1-symmetry EAV DMV pore complex, shown from the x-axis **(a)** and z-axis **(b)**. **(c)** Surface rendering of the C1-symmetry EAV DMV pore complex, shown from the side, top, and bottom. **(d)** Correlation plot (y-axis) between the C1-symmetry EAV pore complex and symmetry orders ranging from C3 to C16 (x-axis). **(e)** Central slice and low-threshold surface render of the C1-symmetry EAV pore complex, highlighting densities associated with the DMV interior (red circle) and the cytosolic side (blue circle) of the channel, respectively. **(f)** Fourier shell correlation (FSC) curves of the C6-symmetry EAV pore complex, corrected FSC (black curve), phase-randomized FSC (red dashed curve) and resolution threshold 0.143 (grey dashed line).

**Supplementary movie 1 – Segmentation model of EAV replication organelles.**

3D segmentation model of the tomogram displaying the following features: the DMV outer membrane (dark yellow), DMV inner membrane (violet), putative RNA filaments (green), pore complexes (coral red), ER (teal green), ribosomes (indigo), nucleocapsids (blue), tubular sheets (Bordeaux red) and rods (purple). Video angles highlight the DMV architecture and association of a spherical nucleocapsid with a pore complex. For visualization, EAV pore complexes and ribosomes were filtered to a resolution of 20 Å.
